# Acetyl-CoA Synthetase 1 regulates global histone propionylation and metabolic stress responses

**DOI:** 10.64898/2026.05.05.722790

**Authors:** Keith C. Garcia, Chengjin Zhu, Ashby J. Morrison

## Abstract

Cells and organisms are often exposed to various metabolic environments that require adaptive responses for survival. One common way cells adapt to fluctuating nutrient environments is through regulated transcription of metabolic genes. Intermediary metabolites, such as acetyl-CoA, produced by metabolic pathways, serve as cofactors for histone post-translational modifications, which in turn regulate gene expression. However, increasing evidence shows that non-acetyl acyl-CoAs, such as propionyl-CoA, participate in gene regulation during metabolic stress. In this report, we find that histone propionylation functions as a global response to glucose starvation. Furthermore, we find that Acetyl-CoA Synthetase 1 (Acs1) binds chromatin and is the primary enzyme responsible for generating propionyl-CoA in the nucleus. Together, our findings reveal that Acs1-mediated histone propionylation constitutes a novel pathway for metabolic adaptation, linking nutrient availability to chromatin modification.

## INTRODUCTION

During times of nutrient stress, cells must respond appropriately to adapt and survive. One common way to adapt to a dynamic nutrient environment is through the regulated transcription of metabolic genes that influence survival and cell growth. An effective way to regulate transcription rapidly and reversibly is through chromatin modification. These modifications require precise coordination between chromatin regulatory machinery and the enzymes responsible for their deposition, many of which depend on intermediary metabolites, such as acetyl-CoA for acetylation and S-adenosyl methionine for methylation of histone tails^1–3^. In nutrient-rich conditions, glucose metabolism promotes acetyl-CoA production through pathways that support biosynthesis and cell growth. In budding yeast, acetyl-CoA synthetase 2, Acs2, is a major contributor to the nuclear acetyl-CoA pool used for histone acetylation under glucose-rich conditions^4^. Consistent with this role, Acs2 is predominantly nuclear and chromatin-associated, suggesting that local acetyl-CoA synthesis may directly support transcriptional programs linked to proliferation^4–7^. Indeed, acetyl-CoA availability promotes histone acetylation at ribosomal protein and ribosome biogenesis genes, linking nutrient abundance to transcriptional programs that support growth^8,9^. A comprehensive understanding of these connections is essential for elucidating the mechanisms that govern cellular and organismal homeostasis.

Advances in mass spectrometry have generated a large catalogue of histone post-translational modifications, raising the possibility that non-acetyl acyl-CoAs may be produced differentially as cells adapt to changing metabolic conditions^10,11^. We have previously shown that as cells transition from gene expression programs associated with pro-growth pathways, such as ribosome biogenesis and translation, to fatty acid β-oxidation pathways, there is a corresponding shift from histone acetylation to histone crotonylation as nutrients become limiting^12^. Moreover, work in mammalian systems has shown that propionyl-CoA, derived from amino acid catabolism and fatty acid oxidation, drives histone propionylation at gene promoters to facilitate transcriptional regulation^13–17^. However, the naturally occurring nutrient environments that promote propionyl-CoA accumulation and the enzymes responsible for its synthesis remain poorly understood.

These observations suggest that distinct metabolic states may use different acyl-CoA-producing enzymes to shape chromatin regulation. While Acs2 has been implicated in acetyl-CoA production and histone acetylation during glucose-rich growth, the enzyme responsible for coordinating acyl-CoA synthesis when glucose becomes limiting has not been established. This raises the possibility that alternative acetyl-CoA synthetase isoforms may support non-acetyl histone acylation during nutrient stress.

In this report, we used a glucose-starvation model in yeast to examine how changes in carbon metabolism alter histone acylation. Utilizing *in vitro* and *in vivo* assays, we identify the acetyl-CoA synthetase isoform, Acs1 as the enzyme responsible for propionyl-CoA synthesis and subsequent histone propionylation across the genome. Collectively, these results expand our understanding of how the metabolic state of the cell connects to genome regulation.

## RESULTS

### Histone propionylation is induced upon glucose starvation and is enriched at promoters genome-wide

To examine histone acylation changes that may underlie chromatin adaptation to carbon stress, cells were grown in glucose-rich media (+dextrose) to mid-log phase and then shifted to glucose-free media (−dextrose) for 16 hours. Histone acetylation, butyrylation, and propionylation were screened via western blot analysis. Consistent with previous studies, histone H3 acetylation was significantly reduced during glucose starvation relative to that of cells in glucose-rich media^15,18^. The abundance of butyrylation exhibited a similar trend to acetylation and was largely reduced during starvation. Interestingly, histone propionylation was the only mark elevated in glucose-depleted conditions (**Figure 1A**).

**Figure 1.**
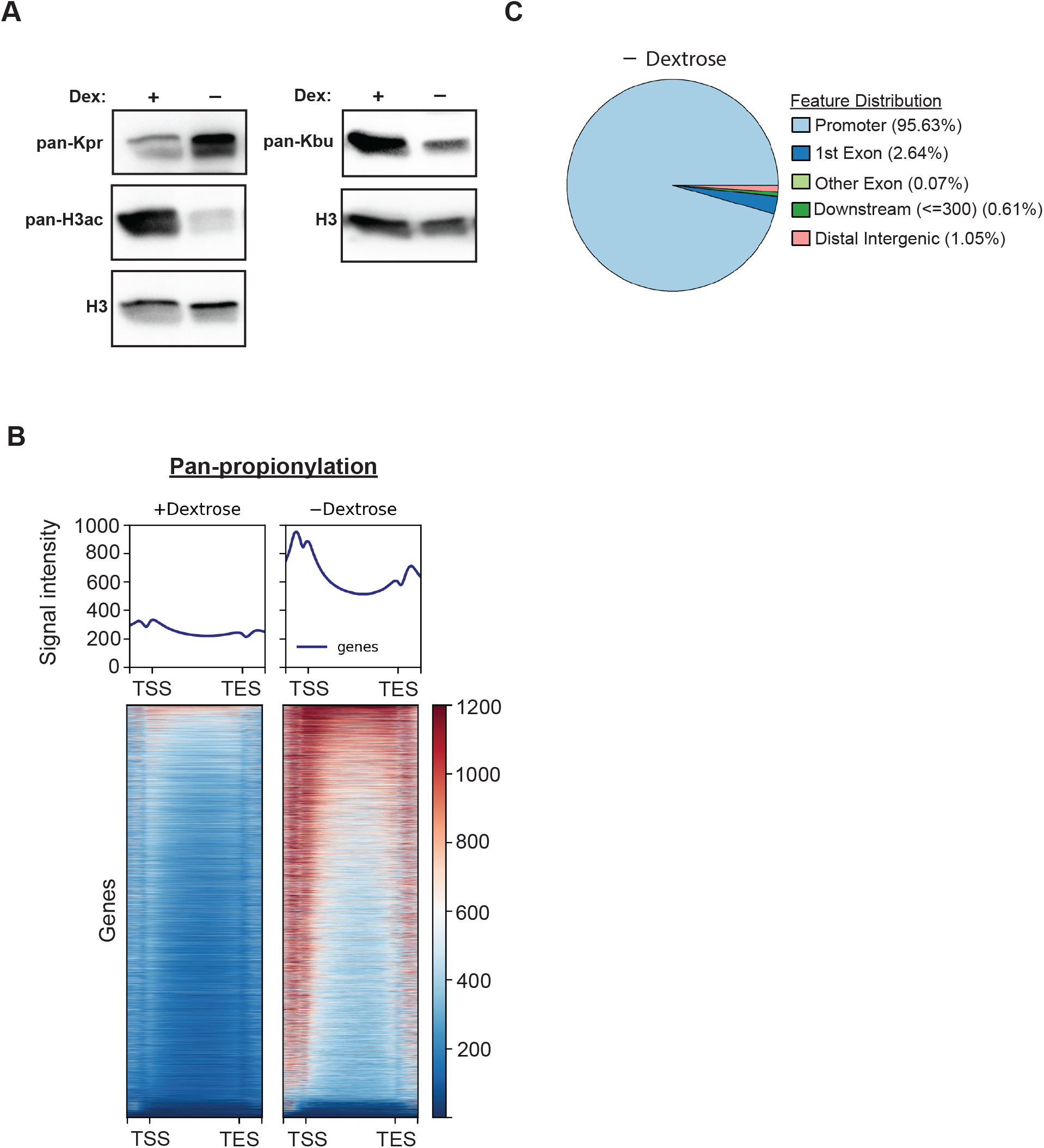
Histone propionylation is elevated during glucose starvation and enriched at promoters. (**A**) Western blots showing differential histone acylation during glucose starvation (16 hours). H3 is used as a loading control. Pan-propionyllysine and H3 acetylation blots were run separately from the pan-butryllysine blot. (**B**) CUT&RUN metagene plots and heatmap showing differential histone propionylation in glucose-rich (+dextrose) and starved (-dextrose) conditions. (**C**) Propionylation peak distribution across features in cells lacking dextrose.

We next performed CUT&RUN analysis to evaluate the effect of glucose starvation on histone propionylation across the genome. In agreement with our western blots, histone propionylation is elevated during glucose starvation (**Figure 1B**). Additionally, CUT&RUN signal was predominantly localized at gene promoters in the absence of glucose (**Figure 1C**). Altogether, these results demonstrate that histone propionylation is a key modification attuned to the metabolic state of the cell, further suggesting a potential role in genome regulation under conditions of glucose scarcity.

### The yeast acetyl-CoA synthase 1 (Acs1) regulates histone propionylation

To identify the enzyme responsible for propionyl-CoA synthesis we focused on Acs1, as prior work implicated it in propionyl-CoA metabolism (**Figure 2A**)^19^. In our yeast starvation model, removal of glucose for 16 hours resulted in elevated Acs1 protein levels, in contrast to Acs2, which remained stable (**Figure 2B**). Since genetic deletion of *ACS2* renders cells inviable when grown on glucose, we utilized a tetracycline (Tet-OFF) inducible system to reduce *ACS2* expression^4^. We optimized doxycycline exposure to a 4-hour treatment, which was sufficient to reduce Acs2 protein levels without affecting cell survival (**Figure S1A** and **S1B**).

**Figure 2.**
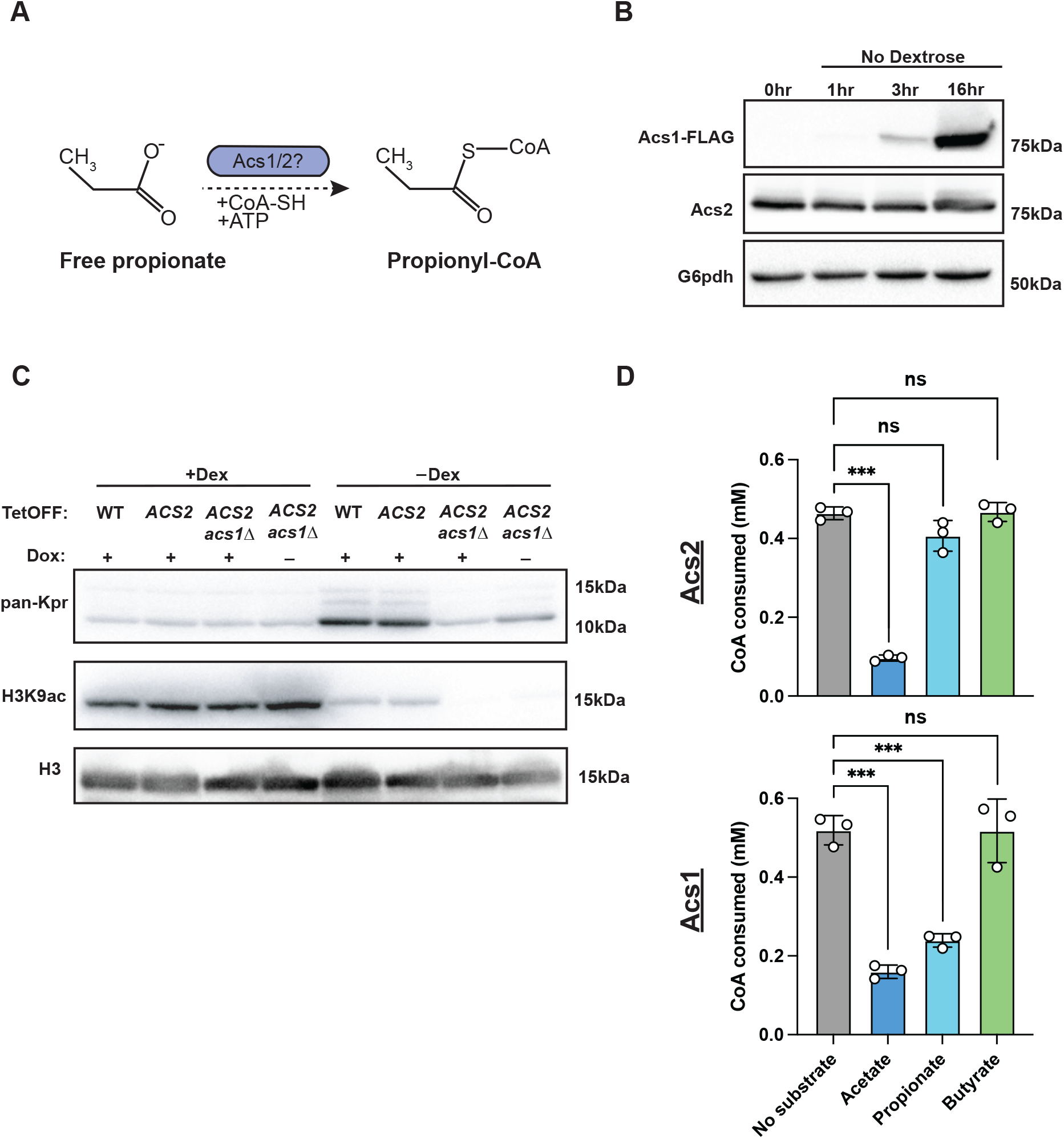
Acetyl-CoA synthetase 1 regulates histone propionylation during glucose starvation. (**A**) Schematic diagram of propionyl-CoA synthesis via Acs1 or Acs2. (**B**) Western blots of Acs1-FLAG and Acs2. Cells were grown to mid-log phase and starved of glucose for 0, 1, 3, and 16 hours. G6pdh is used as the loading control. (**C**) Western blot analysis of acid-extracted histones in the presence or absence of glucose and doxycycline for the indicated Tet-Off strains. Histone propionylation and acetylation are examined using the indicated antibodies. H3 is used as a loading control. (**D**) *In vitro* acyl-CoA synthesis in the presence of CoA with either acetate, propionate, or butyrate. Concentration of CoA consumed compared to no substrate control (free CoA-SH). Statistical significance was assessed using a two-tailed unpaired Student’s *t*-test; ***p<0.001. The data are represented as mean ± SD (n = 3).

Using this system, cells were grown to mid-log in glucose with or without doxycycline, and subsequently shifted to glucose-free media for 16 hours. As before, western blots of histones showed that wild-type cells subjected to starvation exhibited an increase in histone propionylation (**Figure 2C**). Concomitantly, we detected a considerable decrease in H3K9 acetylation, a mark sensitive to glucose withdrawal^15^. Notably, loss of Acs1 results in a significant reduction in histone propionylation during glucose starvation, while the loss of Acs2 does not affect this modification. Combined perturbation of both Acs2 and Acs1 results in a modest additional decrease, suggesting that Acs1 is the predominant isoform responsible for regulating histone propionylation during glucose starvation.

To further examine whether Acs1 is the predominant enzyme responsible for propionyl-CoA synthesis, we performed an *in vitro* acyl-CoA synthesis assay using recombinantly purified Acs1 and Acs2. Acyl-CoA synthetase activity was measured in the presence of a fixed concentration of coenzyme A (CoA-SH) with either acetate, propionate, or butyrate. We quantified the consumption of CoA-SH relative to a no-enzyme control (CoA-SH only) as a functional readout of acyl-CoA synthesis. Reactions in the presence of Acs1 or Acs2 with acetate demonstrate that both enzymes are capable of synthesizing acetyl-CoA *in vitro* (**Figure 2D**). Neither Acs1 nor Acs2 exhibits activity toward the formation of butyryl-CoA. Remarkably, Acs1, but not Acs2, was capable of synthesizing propionyl-CoA *in vitro*. Taken together, these findings identify Acs1 as the predominant isoform responsible for producing cellular propionyl-CoA, which serves as a substrate for histone modification during glucose starvation.

### Acs1 is chromatin-bound during glucose starvation

As previously mentioned, propionyl-CoA can be produced in the mitochondria and peroxisome^13,20,21^. To determine the subcelular localization of Acs1 that is responsible for histone propionylation, we utilized fluorescent microscopy. In glucose-starved cells, we observed that the Acs1 protein was expressed throughout the cell, including the nucleus, where it co-localized with histone H2B (**Figure 3A**). Consistent with previous western results, no Acs1 signal was observed in glucose-rich conditions. In contrast, Acs2 predominantly exhibited a strong nuclear signal in both glucose-rich and glucose-starved media.

**Figure 3.**
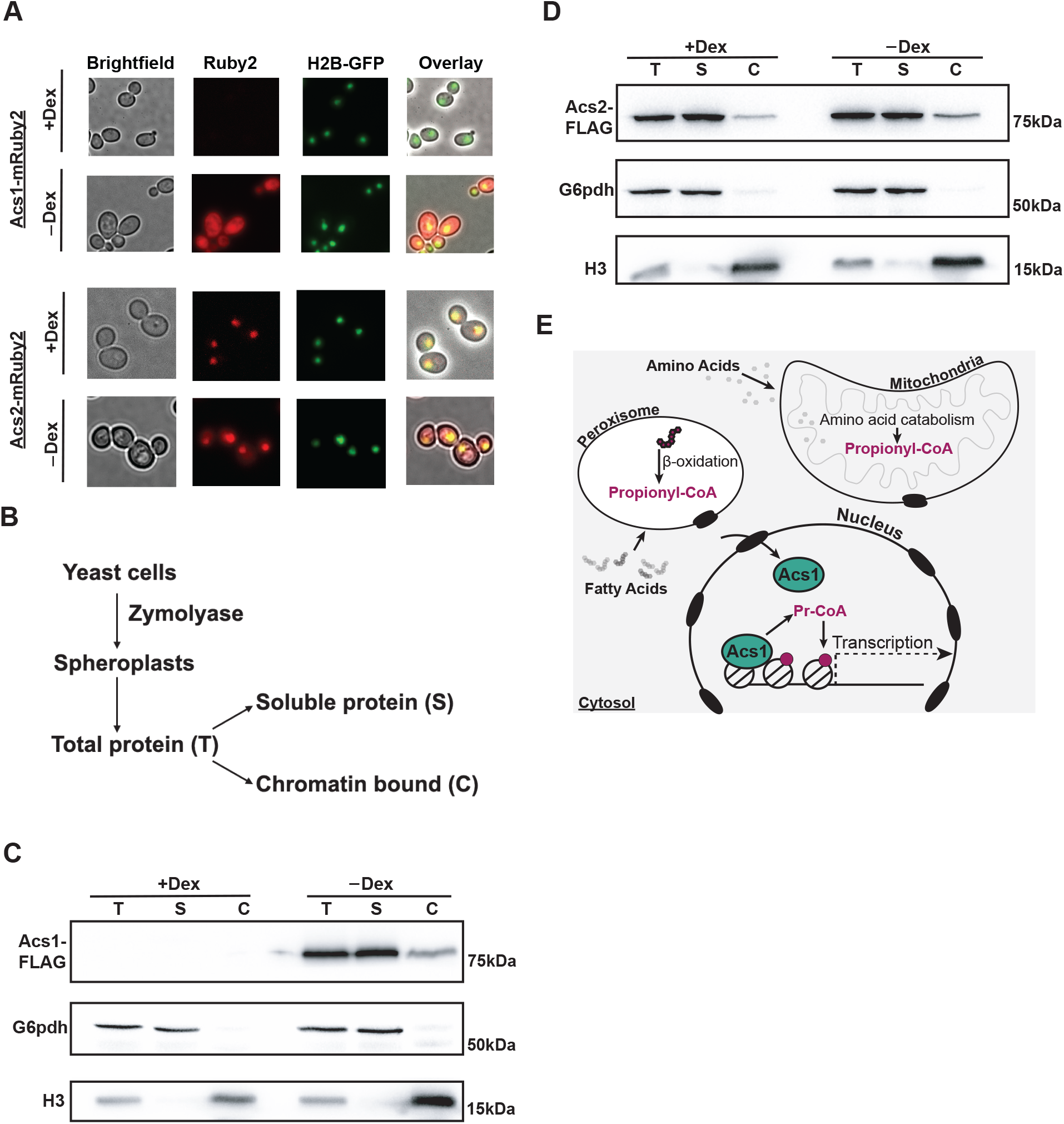
Acs1 is chromatin-associated during glucose starvation. **(A)** Representative z-stack of fluorescence images of Acs1-mRuby2 (top) and Acs2-mRuby2 (bottom) during glucose-rich and starved conditions. H2B-GFP is used as a nuclear marker. (**B**) Schematic of chromatin fractionation protocol. (**C** and **D**) Chromatin fractions for Acs1-FLAG and Acs2-FLAG in glucose-rich and starved conditions. T=total protein, S=soluble fraction, C=chromatin fraction. G6pdh and histone H3 are used as soluble and chromatin fraction controls, respectively. (**E**) A model for Acs1-dependent histone propionylation during glucose starvation. Peroxisomal fatty-acid - oxidation and amino acid catabolism in the mitochondria represent putative sources of propionyl-CoA. Chromatin-associated Acs1 synthesizes nuclear propionyl-CoA for histone propionylation for downstream transcriptional regulation.

Given that Acs1 can localize to the nucleus, we investigated its ability to associate with chromatin using chromatin fractionation assays (**Figure 3B**). In agreement with our fluorescent assays suggesting a nuclear role for Acs1, we observed that Acs1 was enriched in both the soluble and chromatin fractions in cells lacking glucose, whereas Acs2 was observed in all fractions under both glucose-rich and starved conditions. (**Figure 3C** and **D**). Collectively, these results demonstrate that Acs1 is nuclear and chromatin-associated and suggest that Acs1 is responsible for the nuclear pool of propionyl-CoA required for histone propionylation during glucose starvation.

## DISCUSSION

A central outstanding question in chromatin biology is how the metabolic environment connects to histone modification. Specifically, it remains unclear under what conditions many non-acetyl acyl-CoAs accumulate in the cell, and which enzymes are responsible for their nuclear synthesis. Here we show that adaptation to carbon stress elicits a genome-wide switch in histone acylation.

Notably, histone propionylation becomes an abundant modification in glucose-poor environments, while acetylation, the dominant modification in glucose-rich environments, is significantly decreased. Importantly, through *in vitro* and *in vivo* assays, we identify Acs1 as the key acyl-CoA synthetase responsible for nuclear propionyl-CoA synthesis and histone propionylation during glucose starvation (**Figure 3E**).

### A nuclear role for Acs1

Pioneering studies in yeast identified a nuclear function for Acs2 in the regulation of histone acetylation, and work in mammalian cells has shown that the homolog ACSS2, together with ATP-citrate lyase (ACLY), synthesizes nuclear acetyl-CoA for chromatin modification^4,22–24^. Our results show that, like Acs2, Acs1 is nuclear and chromatin-associated. Our studies further support the hypothesis that, under glucose-limiting conditions, Acs1 is the predominant isoform responsible for nuclear propionyl-CoA synthesis and its subsequent deposition onto histones. However, it remains to be understood how Acs1 is localized into the nucleus and which genomic regions it occupies. In contrast, Acs2 has been shown to occupy subtelomeric regions in glucose-rich conditions, where it maintains a pool of H4K16 acetylation to facilitate telomere silencing^6^. In light of these findings, an important open question is whether Acs1 functionally recapitulates a similar preference for binding chromosomal ends during glucose starvation and whether it has additional roles in chromatin and gene regulation.

### Propionyl-CoA metabolism in yeast

As mentioned, the modification of histones is dependent on intermediary metabolites, which are generated from varying metabolic processes and are produced in different cellular compartments^13,14,20,25^. During glucose fermentation in the yeast cytosol, glucose is converted to pyruvate via glycolysis. Pyruvate is decarboxylated to acetaldehyde, which is reduced to ethanol and acetate, which is converted to acetyl-CoA^26,27^. Under these glucose-rich conditions, histone acetylation is abundant, and cells respond by expressing genes involved in cell growth (e.g., ribosomal proteins)^8,12^. Conversely, when glucose becomes limiting, cells reprogram their gene expression to upregulate fatty acid degradation enzymes, thereby promoting the production of acetyl-CoA and other acyl-CoAs within peroxisomes^20,21^. It has been observed that additional sources of propionyl-CoA arise from the mitochondrial degradation of branched-chain amino acids (valine, leucine, and isoleucine), as well as from methionine and threonine catabolism^13,14,28^. However, these pathways have been primarily characterized in mammalian systems, and whether they similarly contribute to propionyl-CoA in yeast remains unclear. It is important to note that the presence of propionate in the cytosolic pool indicates that propionyl-CoA in these compartments undergoes transesterification back to propionate for export^29,30^. Free propionate can then be imported into the nucleus, where Acs1 reconverts it to propionyl-CoA for histone modification. This nuclear propionate to propionyl-CoA conversion represents a potential mechanism coupling cytosolic metabolic flux to chromatin modification.

### The role of histone propionylation in transcriptional regulation

Prior work in mammalian cells has provided evidence that histone propionylation is emerging as a key modification involved in transcriptional regulation^16^. Studies by Nitsch et al. link elevated levels of histone propionylation in mouse models of propionic acidemia, a metabolic disorder resulting in the accumulation of propionic acid in the liver^17^. This study and work by others have shown that the bulk of histone propionylation is enriched at promoters and linked to transcriptional activation^17,31^. Other studies in colon cancer cells have shown that exogenous propionate increases histone propionylation at promoters, which is associated with greater chromatin accessibility and subsequent upregulation of proto-oncogenes involved in cellular proliferation^32^. Here, we show that glucose starvation in yeast leads to a genome-wide increase in histone propionylation. Notably, this modification is mostly enriched at transcription start sites, suggesting a role in transcriptional regulation during glucose stress and indicating that this mark may have a functionally conserved role across different biological contexts. Additional studies are warranted to determine whether propionylation directly regulates metabolic genes required for adaptive cellular responses during glucose starvation.

### Broader implications of histone propionylation in disease

To date, histone propionylation has been observed across different disease contexts, including cardiovascular disease, neurological disorders, and cancer^28,33–35^. Thus, the link between metabolic state and chromatin modification is becoming increasingly recognized as a critical determinant of human health, shaping gene expression programs that underlie both normal physiology and pathology. Additionally, understanding the enzymes responsible for synthesizing these substrates and the diverse contexts in which their functions are co-opted in disease is critical for future therapeutic opportunities. Overall, this study demonstrates that the nuclear function of metabolic enzymes plays a context-dependent role in coupling nutrient availability to chromatin regulation, thereby driving adaptive responses to nutrient stress with broad implications for human health and disease.

## Methods

### Yeast strains and growth conditions

The *Saccharomyces cerevisiae* strains used in this study are listed in **Supplementary Table S1**. All strains are derived from the S288C background. All strains were cultured at 30°C. Glucose-rich cells were grown in YPD to an OD660 of 0.75-0.8. For glucose starvation, strains were grown to an OD660 of 0.75-0.8 in YPD, spun down and washed in sterile dH2O, and transferred to YP media lacking dextrose for 16 hours.

Tet-Off strains were taken from stationary phase and inoculated to an OD 0.3 into fresh YPD. For the 4-hour treatment, cells were grown for 1 hour prior to doxycycline addition (50 μg/ml), and samples were collected. For the 5-hour treatment, cells were inoculated with doxycycline (50ug/ml) for 5 hours before sample collection. For glucose starvation, strains were prepped as mentioned above, spun down and washed in sterile dH2O, added with to YP media with fresh doxycycline (50ug/ml), and incubated overnight for 16 hours before sample collection.

### Recombinant Acs1 and Acs2 purification

*Saccharomyces cerevisiae ACS1* and *ACS2* genes were cloned into the pETM40 expression vector with a C-terminal TEV-10XHis tag. Expression vectors were transformed into BL21 for recombinant protein expression. 1 liter LB+Kanamycin culture was inoculated at a starting OD600 of 0.1 and allowed to grow at 37°C at 180rpm to an OD600 of 0.6. The cultures were removed and allowed to cool to room temperature, after which 1mM IPTG was added, and cells were grown overnight at 18 °C at 180rpm. Cells were harvested at 4000rpm at 10 °C (JLA 8.1000 Beckman rotor). The pellet was resuspended in lysis buffer (80mM HEPES pH 7.4, 200mM NaCl, 0.5mM MgCl2, 10%glycerol) supplemented with 0.2mM PMSF, 1mM benzamidine, 1mM DTT, 1mM β-mercaptoethanol, protease inhibitors, DNase, and lysozyme. Bacterial resuspension was lysed using a microfluidizer at 18,000psi. Lysate was clarified by ultracentrifugation using a Ti45 rotor at 10,000rpm for 1 hour at 4 °C.

The supernatant containing Acs1 and Acs2 proteins were first purified by passing over a 5 ml HisTrap HP column (Cytiva) and washed with 70mM imidazole, then eluted at 300 mM imidazole. The eluted proteins were then loaded onto a Hiload 16/600 Superdex 200 pg (Cytiva). The column was equilibrated with GF buffer (10 mM HEPES pH RT 7.4, 200 mM NaCl, 0.5 mM MgCl2, 5% Glycerol, 1 mM DTT). Peak fractions were collected, flash frozen, and stored at -80 °C. The final proteins were concentrated to 1.5 mg/ml, aliquots were flash frozen, and stored at −80 °C.

### Western blotting and chromatin fractionation assay

10mL of yeast was grown to an OD660 of ∼0.75-0.8. Cell pellets were resuspended on ice with 300 μL of cold TCA lysis Buffer (10% trichloroacetic acid, 10mM Tris-HCl, pH 8.0, 25mM ammonium acetate, and 1mM EDTA, pH 8.0). Glass beads (approximately half the volume) were added to the cellular suspension and were subsequently lysed using the bead-beating method on a FastPrep-24 (MP Biomedicals) at 6.0 m/s for 1min repeat four times at 4°C. Crude lysate was centrifuged for 10 minutes at 16,000 x g, at 4 °C. The precipitated protein and cellular debris were resuspended in 150 μL of resuspension buffer (0.1M Tris-HCl, pH 11.0, and 3% SDS). Samples were then boiled at 100 °C for 5 minutes and centrifuged at 16,000 x g for 30 seconds. The supernatant was collected, and protein quantification was performed by BCA assay. Proteins were detected by western blot using anti-FLAG (Sigma M2; F1804), anti-H3 (Epicypher; 13-0001), anti-Glucose-6-phosphate dehydrogenase (G6pdh) (Millipore/Sigma; A9521), anti-propionyllysine (PTM BIO; PTM-201), and anti-butryllysine (PTM BIO; PTM-301).

Chromatin fractions were performed as previously described^36^ with some modifications. 3×10^8^ cells were spheroplasted with zymolyase 100T at 30 °C for 15-20 minutes for cells in YPD, and 1.5-2 hours for glucose-starved cells. Spheroplasting was examined both spectrophotometrically and microscopically. For the insoluble white pellet, 200 μL of lysis buffer was added to resuspend the pellet. Approximately 20 μL of glass beads was added to each sample and vortexed at 4 °C for 5 minutes. Anti-G6pdh and H3 were used as cytoplasmic and nuclear fraction controls, respectively.

### Immunofluorescence

For colocalization with the fluorescently tagged strains, cells were gently fixed for 10 minutes with 3% formaldehyde before imaging. Z-stack images were collected on a Leica DMi8 fluorescent microscope.

### DTNB (5,5-dithiobis-(2-nitrobenzoic acid) assay

3uM of recombinant Acs1 or Acs2 was incubated for 10 minutes at 30 °C with 4mM ATP, 0.5mM coenzyme A, 0.5mM sodium acetate, sodium propionate, or sodium butyrate in a 25uL reaction volume containing reaction buffer (0.1M Tris-HCl, pH 8.0, 5mM MgCl2). Enzyme reactions, along with a standard curve of coenzyme A, were quickly added to 225 μL reaction buffer containing 0.2 M DTNB. Absorbance was measured at 412nm on a Synergy microplate reader at room temperature. Interpolated 412nm readings from the standard curve were used to determine the concentration of coenzyme A consumed in the enzymatic reactions.

### CUT&RUN sample preparation

Crude yeast nuclei were prepared as described with some modifications^37^. SPC buffer (1M sorbitol, 200mM PIPES, pH 6.3, 0.1mM CaCl2) was supplemented with yeast protease and histone deacetylase inhibitors (5mM sodium butyrate and 250nM trichostatin A). Cells were grown to an OD of 0.75-0.8. Before sample collection, cells were crosslinked (0.5% formaldehyde) for 1 minute and quenched with a 0.125M final concentration of glycine. Approximately 3 × 10^8^ cells were collected and spheroplasted with zymolyase 100T at 30 °C for 15-20 minutes for glucose-grown cells and 1.5-2 hours for overnight glucose-starved cells. Spheroplasts were examined both spectrophotometrically and microscopically. After Ficoll nuclei isolation, nuclei were aliquoted into 8-10 tubes with approximately 2 × 10^7^ nuclei, placed in an isopropyl chamber for slow freezing at -80 °C.

Yeast CUT&RUN was performed as described with a few modifications^37^. Approximately 1.5 × 10^7^ crude nuclei were used per reaction. Frozen nuclei were placed on a 37 °C heat block just until the sample began to thaw. Tubes were quickly placed on ice to allow the rest of the sample to thaw completely. 40 μL of activated concavalin A beads were added to each reaction and rotated end over end for 1.5 hours at room temperature. After binding, samples were placed on a magnetic rack and resuspended in 200 uL of digitonin-wash buffer. Anti-pan-propionyllysine antibody was added at a 1:100 dilution, and isotype negative control IgG at a concentration of 200 μL in a reaction. Samples were incubated overnight on a nutator at 4 °C. The following day, samples were washed twice and resuspended in 200 μL. 5 μL of pAG-MNase was added to each sample and incubated at 4 °C for 1 hour on a nutator. Before digestion, samples were washed three times, resuspended in 200 μL of digitonin-wash buffer, and kept on an ice-water bath for 10 minutes. 2mM final concentration of CaCl2 was added simultaneously while the samples were gently vortexed. After the addition of CaCl2, samples were immediately placed on ice, and the digest was allowed to proceed. After digestion, 200 μL of 2xSTOP buffer containing *E*.*coli* spike-in was added. Samples were then incubated for 20 minutes at 37 °C to release CUT&RUN fragments. Fragments were subsequently de-crosslinked (0.01% SDS and 0.2mg/ml proteinase K) at 55 °C overnight. DNA purification was performed using a 1.8X bead concentration. Samples were eluted in 55 μL of 0.1X TE buffer.

### CUT&RUN library preparation and sequencing

Libraries were prepped from purified CUT&RUN DNA fragments (3-4ng input) using the NEBNext Ultra II DNA Library Prep Kit (E7645S) for Illumina with some modifications. End Prep dA tailing reactions were incubated for 30 minutes at 20 °C and 1 hour at 50 °C (this lower temperature prevents smaller DNA fragments from denaturing but is still able to heat-inactivate the enzyme)^38^. NEBNext Unique Molecular Identifier (UMI) adapters (E7878S) were diluted 1:50 for adaptor ligation reactions, followed by clean-up of adaptor-ligated DNA without size selection. PCR enrichment (15 cycles) of adapter-ligated DNA used an annealing/extension time of 10 seconds at 65 °C. Following adapter clean-up, the quality of libraries was assessed on an Agilent 4200 TapeStation. Before sequencing, libraries were quantified using the NEBNext Library Quant Kit (E7630S) for Illumina. Libraries were sequenced on a Novaseq X plus (2×150).

### CUT&RUN data processing

Data processing was performed as previously described^39^. Paired-end fragments were aligned to the sacCer3 reference genome (bowtie)^40^. UMIs were extracted and used to remove PCR duplicates (UMI-tools). A spike-in scaling factor was calculated for each library (samtools idxstats) and applied to generate normalized bigwig files (bedGraphToBigWig)^41,42^. Heatmaps of spike-in normalized CUT&RUN signal were generated using deepTools^43^. Peak annotation and genomic features were determined using ChIPseeker^44^

### Acid extraction of yeast histones

Crude yeast nuclei were prepared as described previously and subsequently lysed in high salt extraction buffer (10mM MES, pH 6.0, 430mM NaCl, and 0.5% NP-40) supplemented with yeast protease inhibitors, 5mM NaButyrate, and 250nM trichostatin A. Cells were placed on ice for 5min, spun down at 13000 rpm, and supernatant removed. The pellet was washed twice in high salt extraction buffer. To the crude chromatin pellet, 5 packed volumes of 0.2M H2SO4 were added, and the pellet was resuspended completely. Samples were rotated at 4 °C for 2 hours and spun down at 13000 rpm for 15 minutes at 4 °C. Supernatant was collected, and 8 volumes of cold acetone were added to the sample, and histones were precipitated overnight at -20 °C. Precipitated histone/protein was spun down at 3500 rpm for 10 minutes, washed twice in cold acetone (centrifuged for 5 minutes at 10000 rpm). Residual acetone was evaporated/removed on a 37 °C heat block, and the pellet was resuspended in 60 μL of Mass Spectrometry-grade water.

## ACKNOWLEDGMENTS

We thank members of the Morrison Lab for their suggestions and helpful discussions. We thank Dr. Naima Sharaf for help with Acs1 and Acs2 purifications. We also thank Jeff Boeke for the Acs2 antibody.

## AUTHOR CONTRIBUTIONS

Conceptualization, K.C.G. and A.J.M.; methodology, K.C.G. and A.J.M. Investigation, K.C.G., C.Z., and A.J.M; writing—original draft, K.C.G. and A.J.M.; writing—review & editing, K.C.G. and A.J.M.; funding acquisition, K.C.G. and A.J.M.; resources, A.J.M.; supervision, A.J.M.

