## Supplementary Figure for "Acetyl-CoA Synthetase 1 regulates global histone propionylation and metabolic stress responses"

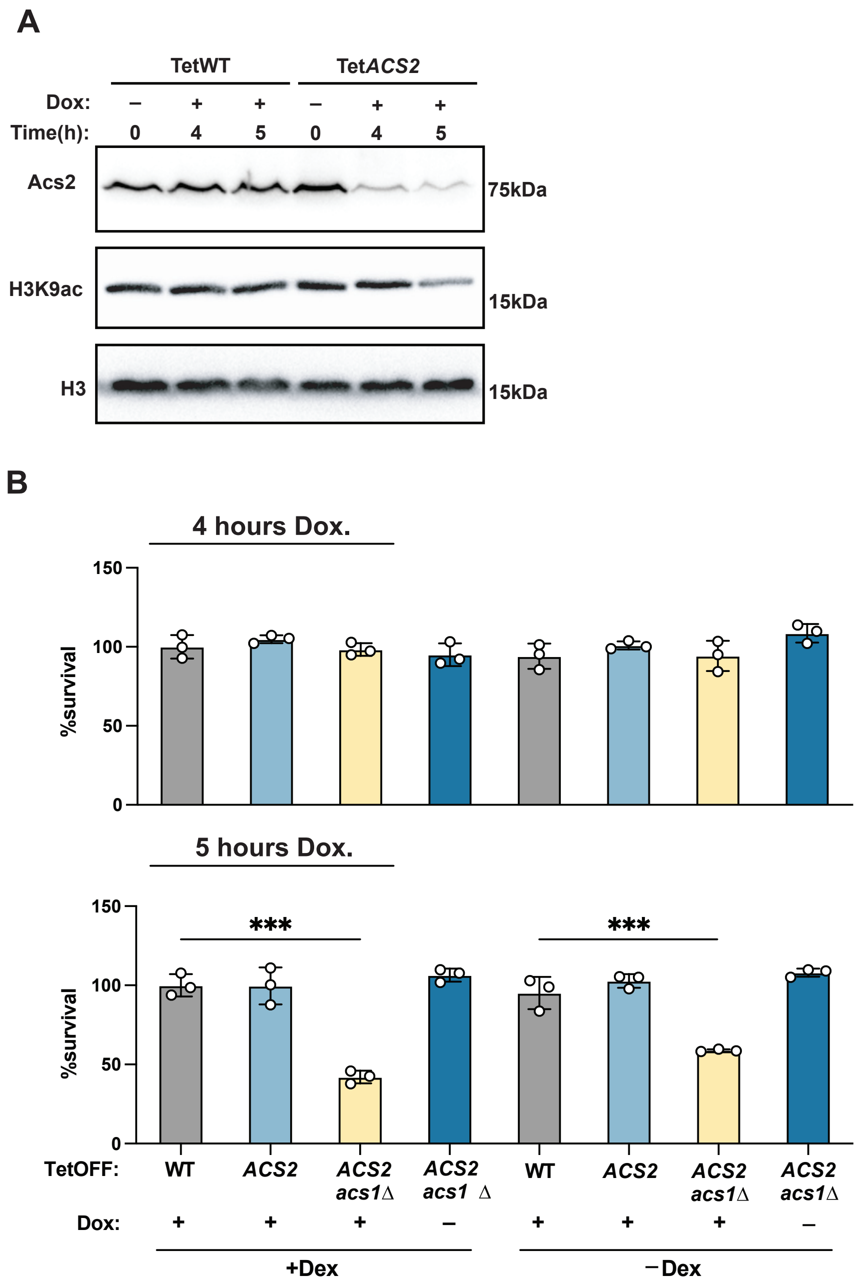


**Figure S1. Differential expression of Acs1 during metabolic stress and the impact of Acs2 loss on cell survival. Related to Figure 2.** (**A**) Western blot with indicated antibodies of a time course for doxycycline-treated (50ug/mL) TetOFF-WT and TetOFF-*ACS2* strains. Histone H3 is used as a loading control. (**B**) Cell survival of Tet-strains exposed to doxycycline for 4 (top) or 5 hours (bottom) in YPD prior to being shifted to glucose free media. Statistical significance was assessed using a two-tailed unpaired Student's *t*-test; ***p<0.001. The data are represented as mean ± SD (n = 3).
