## Supplementary Table for "Acetyl-CoA Synthetase 1 regulates global histone propionylation and metabolic stress responses"

| Table S1. Strains and plasmids used in this study. | | |
|---|---|---|
| **Strain** | **Genotype** | **Source** |
| BY4741 | *MATa his3Δ1 leu2Δ0 met15Δ0 ura3Δ0* | Open Biosystems |
| KGY073 | BY4741 *acs1::KANMX4* | Open Biosystems KO collection. Winzeler et al.[45] |
| KGY064 | BY4741 *ACS1-2XFLAG-KANMX6* | This paper |
| KGY082 | BY4741 *ACS2-2XFLAG-KANMX6* | This paper |
| HTB2-GFP | BY4741 *HTB2-GFP-HIS3MX6* | Huh et al[46] |
| KGY085 | BY4741 *ACS2-yomRUBY2-KanR HTB2-GFP-HIS3MX6* | This paper |
| KGY076 | BY4741 *ACS1-yomRUBY2-KanR HTB2-GFP-HIS3MX6* | This paper |
| R1158 | BY4741 *URA3::CMV-tTA* | Hughs et al.[47] |
| Tet-*ACS2* | R1158 *prACS2::kanR-tet07-TATA* | Hughs et al.[47] |
| KGY083 | Tet-*ACS2 acs1::hphMX6* | This paper |
| **Plasmids** | **Description** | **Source** |
| | pETM-40 *ACS1_TEV_10xHis* | This paper |
| | pETM-40 *ACS2_TEV_10xHis* | This paper |
